# Predatory aggression evolved through adaptations to noradrenergic circuits

**DOI:** 10.1101/2024.08.02.606321

**Authors:** Güniz Goze Eren, Leonard Böger, Marianne Roca, Fumie Hiramatsu, Jun Liu, Luis Alvarez, Desiree Goetting, Nurit Zorn, Ziduan Han, Misako Okumura, Monika Scholz, James W. Lightfoot

## Abstract

Behaviors are adaptive traits evolving through natural selection. Crucially, the genetic, molecular, and neural modifications that shape behavioral innovations are poorly understood. Here, we identify specialized adaptations linked to the evolution of aggression in the predatory nematode *Pristionchus pacificus*. Using machine learning, we identified robust behavioral states associated with aggressive episodes. These depend on modifications to the invertebrate noradrenergic pathway, with octopamine promoting predatory bouts, and tyramine antagonistically suppressing predation. Strikingly, aggression coincides with rewiring of key circuits across nematode evolution. We find additional octopaminergic neurons with morphological adaptations, including neurites extending to teeth-like structures, and expanded receptor expression throughout head sensory neurons gating prey detection. Thus, evolutionary adaptations in noradrenergic circuits facilitated the emergence of aggressive behavioral states associated with complex predatory traits.

---

Natural selection favors behavioral traits that enhance an organism’s fitness. This process results in a wide range of behaviors adapted to specific ecological niches and facilitates the evolution of behavioral diversity across species (*1*). Certain behaviors such as foraging, mating, predator avoidance and aggression are subject to strong environmental pressures and are therefore particularly susceptible to evolutionary change. Rare insights into these evolutionary processes have been gleaned from interspecies comparative studies. These include comparisons of *Peromyscus* mice which have evolved distinct burrowing behaviors for predator avoidance and parental care (*2*, *3*), as well as behavioral differences between diverse *Drosophila* species which influence courtship signals and food foraging preferences (*4–7*). Despite this, the precise genetic, molecular, and neural basis of behavioral adaptations are poorly understood.

Nematodes with their small nervous systems and well-developed genetic and molecular tools are potent systems for understanding evolutionary behavioral processes in detail. Specifically, compared to *Caenorhabditis elegans*, the predatory nematode *Pristionchus pacificus* has evolved striking behavioral differences. This includes diversification in odorant sensitivity (*8*), social preference (*9*, *10*), and aggressive behaviors used to establish territory and remove competitors from their environment through intraguild predation and cannibalism (*11–14*). Consequently, we have leveraged the behavioral diversity between these nematode species to investigate the evolutionary adaptations molding the *P. pacificus* aggressive traits.

## Results

*P. pacificus* is an omnivorous nematode species which, in addition to feeding on bacteria, also preys on the larvae of other nematodes (Fig. 1A and Movie S1). Moreover, while predation is utilized by *P. pacificus* to generate an additional nutrient source, it is also an aggressive mechanism for establishing territory through the removal of competitors. This includes the killing of other species and the cannibalism of conspecifics (*11–14*). The increased feeding complexity is associated with the evolution of specialized morphological adaptations and behavioral actions. Specifically, during predation distinct pharyngeal dynamics and the presence of teeth-like armaments facilitate the puncturing of the prey cuticle and feeding on their innards (*15–17*). However, not every contact with potential prey results in an attack, indicating that *P. pacificus* predatory aggression is not indiscriminate (Movie S1 and Movie S2). Therefore, to investigate the mechanism underlying the evolution of aggression and its regulation in *P. pacificus*, we have developed a high-throughput analysis tool to automatically characterize *P. pacificus* feeding behaviors.

**Fig. 1.**
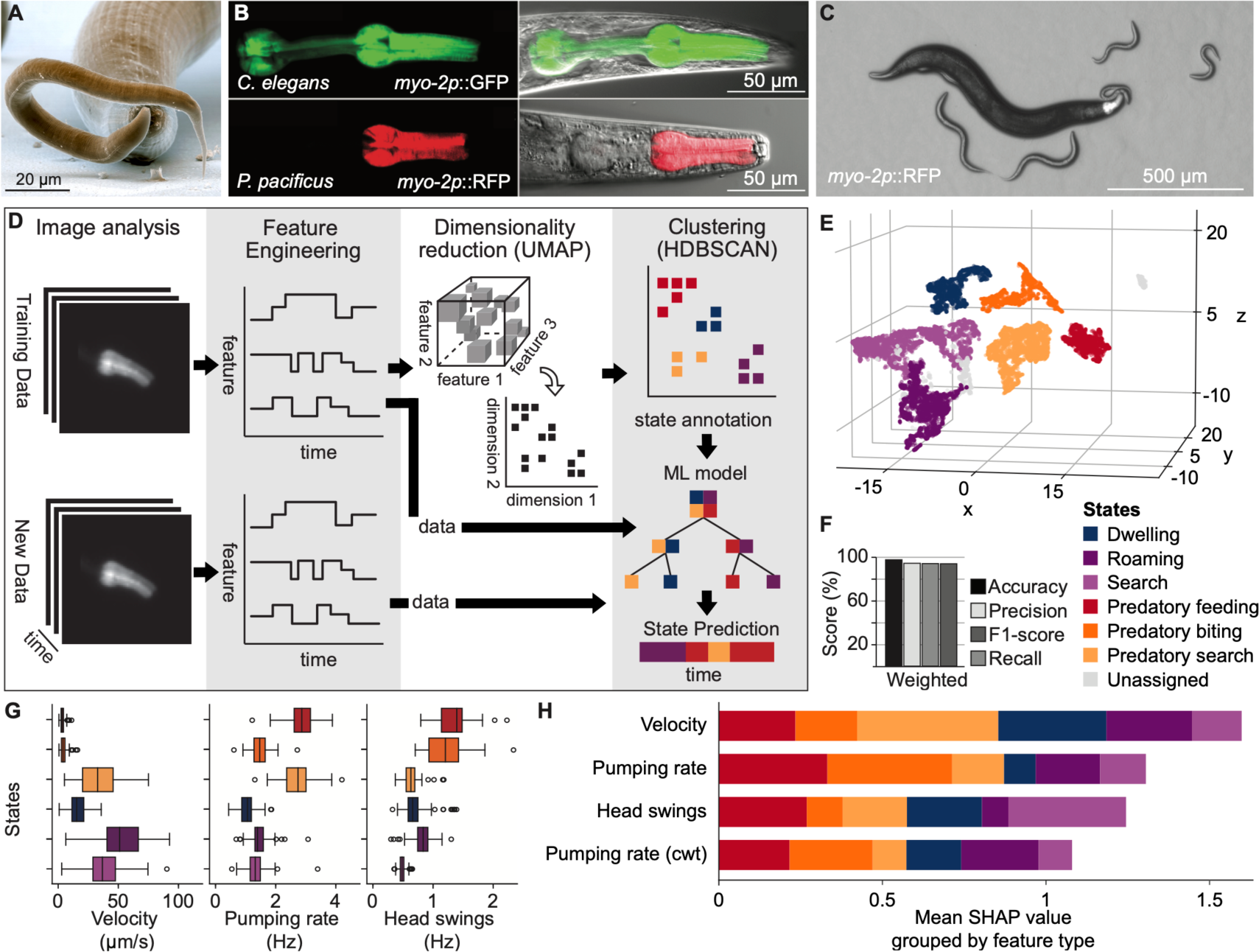
Machine-learning model predicts behavioral states from high-throughput tracking data of a predatory nematode. (**A**) SEM image of *P. pacificus* (background nematode) with *C. elegans* larvae (foreground nematode). (**B**) *myo-2p*::GPF expression in *C. elegans* compared to *myo-2p*::RFP expression in *P. pacificus*. (**C**) Predatory *P. pacificus* animal surrounded by larval *C. elegans* prey. (**D**) Schematic of the machine-learning pipeline used to classify behavioral states. (**E**) UMAP embedding of behavioral features. Colors indicate the six behavioral states identified by hierarchical clustering. (**F**) Performance metrics of the behavioral state classifier on novel, unseen data. (**G**) Distribution of key behavioral features in each state. Each point in the box plots corresponds to the mean value per state and per tracked animal. (**H**) Feature importance (SHAP values) for each of the basic behavioral features involved in the model. Features derived from the same base feature are averaged. Colors correspond to the behavioral states represented in (E).

### Behavioral tracking and state prediction

Nematodes feed via their pharynx, a neuromuscular organ in the animal head. In *C. elegans*, a fluorophore label targeted to the pharyngeal muscle has facilitated the dissection of its feeding behaviors across development and under different environmental conditions (*18*). To track feeding behaviors in *P. pacificus,* we exploited a similar pharyngeal fluorophore label using the *Ppa-myo-2* promoter (Fig. 1B).

This plasmid was integrated into the *P. pacificus* genome with no adverse behavioral effects (fig. S1). We observed much lower levels of *Ppa-myo-2* expression in the terminal bulb of the *P. pacificus* pharynx than in *C. elegans* due to the enlarged gland cells found in this space along with the absence of a hardened grinder structure in *Diplogastridae* (*19*, *20*). Despite this difference, we were able to use this method to successfully track feeding and locomotion in many animals simultaneously on standard assay conditions (Fig. 1C). These consist of *Ppa-myo-2p*::RFP expressing *P. pacificus* predators placed onto an assay arena that contains either *C. elegans* larvae in abundance as potential prey or a bacterial lawn as food source. During behavioral tracking, we extracted multiple features including speed, reversals, feeding events, as well as posture-related measures using the image analysis tool PharaGlow. To automatically identify predatory states, we employed a machine-learning pipeline combining low-dimensional embedding and hierarchical clustering (Fig. 1D). Similar pipelines have been used for unsupervised state classification of behavioral tracking data in flies, mice, fish, and *C. elegans* (*21–27*). Using this approach, we found six distinct behavioral states (Fig. 1E and fig. S2 A, B). We subsequently identified the behaviors in each state based on the overlap with annotations of an expert human annotator (fig. S2C).

Several of these appear to correspond to canonical states also observed in *C. elegans* including ‘roaming’ and ‘dwelling’ states, however, we also identified three novel behavioral states which correlate to a predatory environment. These states are labeled ‘predatory search’, ‘predatory biting’ and ‘predatory feeding’. Next, to extend the model to unseen data, we fit an XGBoost multi-class classifier (*28*) to the clustered data, and analyzed its performance on a test set of held-out recordings. The model captures the cluster labels with > 95% in both accuracy and recall (Fig.1F and fig. S3) with velocity, pumping rate, and head swings contributing most to state identity (Fig. 1 G and H). Consequently, the resulting pipeline allowed us to predict the behavioral states for unrestrained animals (Movie S3), thus providing a mechanism to investigate the molecular determinants driving the evolution of aggression in *P. pacificus*.

### Environmental and sensory context shapes predatory drive

To determine the specificity of our identified behavioral states, we investigated the influence of the environmental and sensory context on *P. pacificus* state occupancy by analyzing ∼ 10.5 animal hours of tracking data on either a bacterial food source or larval prey. The most descriptive features for each state are velocity and pumping rate of the animals (Fig. 1H and fig. S3D). We found that the joint distribution of these two features showed distinct clusters when animals were placed on prey larvae or on bacterial food (Fig. 2A and B). By comparing the density plots with the identified behavioral states, we observed that animals preferentially occupy predation-related states when placed on larvae (Fig. 2B) whereas they spend more time in low pumping rate states with higher speeds when exposed to bacteria (Fig. 2B, right). We identified two types of search states, which we label as ‘search’ and ‘predatory search’. Both are characterized by higher speeds, however, on prey, they showed exaggerated head swing amplitudes during ‘predatory search’ making this state distinct from the ‘search’ state, which is more prominent on bacterial foods (Fig. 2C and D). These findings suggest that sensory context dictates state occupancy such that predatory states are nearly unique to prey-rich environments, further validating our previous annotations (Fig. 2C to F and fig. S2C). Interestingly, we found that while transitions between states and the total time spent in each state change are dependent on sensory contexts, the average duration of a behavioral state remained constant for the majority of states (Fig. 2E and F, and fig. S4A to C). We therefore focused on the total time spent in each state as a measure of the propensity for predatory versus non-predatory behavior. During ‘predatory biting’, *P. pacificus* attacks and punctures the cuticle of their prey with their teeth-like denticles. Therefore, ‘predatory biting’ in *P. pacificus* serves as a robust indicator of aggressive actions and consists of a combination of nutritional drive (‘predatory biting’ to ‘predatory feeding’ (fig. S4D)) and other aggressive interactions such as territoriality and surplus killing (*11*, *12*). Thus, we observe a greatly expanded state complexity in *P. pacificus* including aggressive behaviors compared to those reported in *C. elegans*. This complexity may be a feature of the *P. pacificus* omnivorous lifecycle and dietary switching, or instead a characteristic of the evolution of predation. Future work on obligate predators will elucidate this aspect of state modulation.

**Fig. 2.**
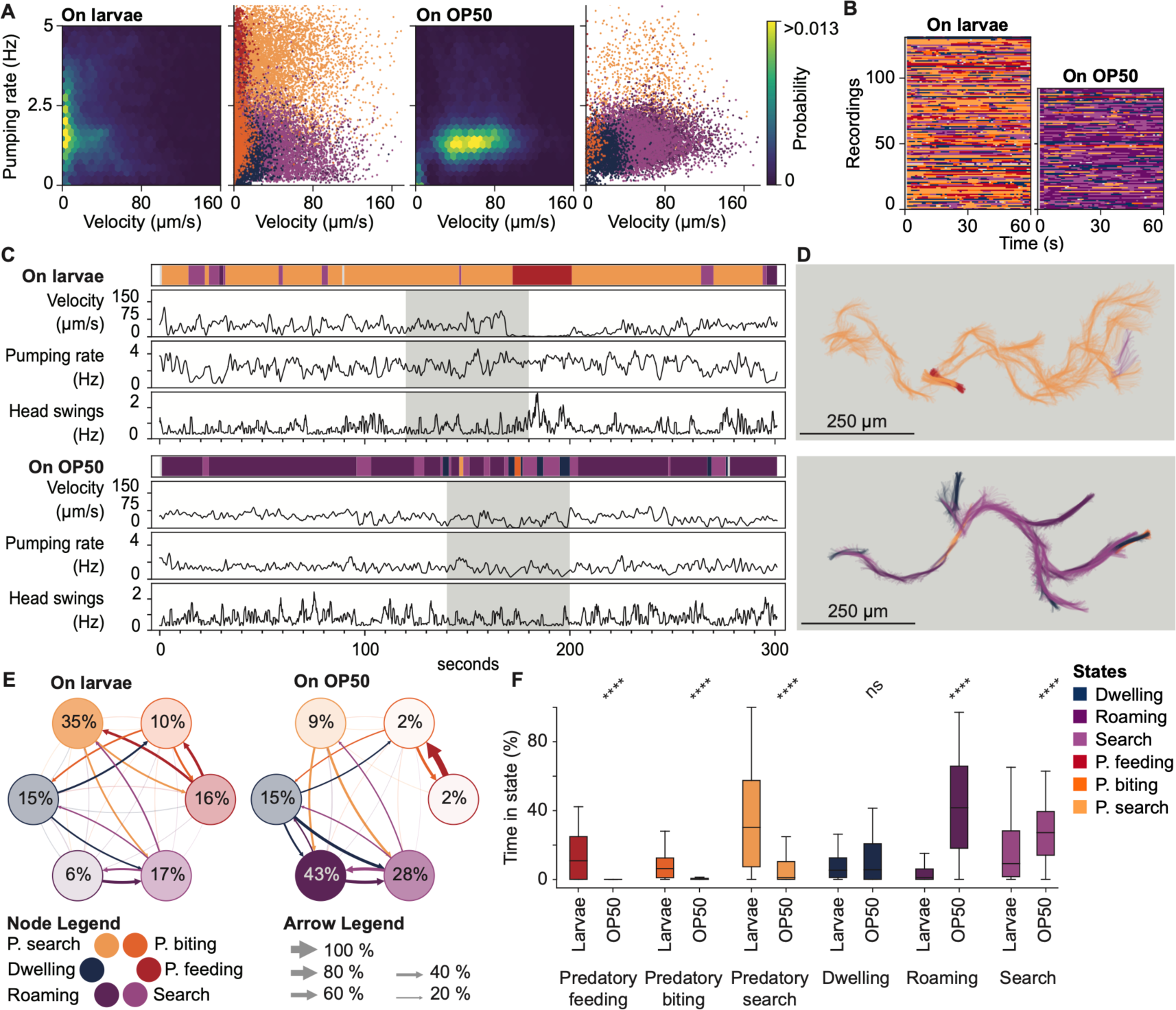
Automatic classification of behavioral data reveals context-dependent predation drive. (**A**) Probability density map of velocity and pumping rate for animals on larval prey or OP50 bacteria. Scatter plots indicate the corresponding state assignments. (**B**) Ethograms showing predicted behavioral states for animals on prey (left) or bacteria (right). (**C**) Ethograms, velocity, pumping rate, and head swings for a representative animal on larval prey (top) or bacterial food (bottom). (**D**) Pharyngeal centerline as colored by the assigned behavioral state. The track shown corresponds to the grey region in (C). (**E**) Average transition rates between behavioral states for animals on larval prey and bacterial food, respectively. Numbers in circles indicate the fraction of time per state as in (E). Arrow thickness indicates the transition rate normalized to out-going transitions. (**F**) Mean fraction of time spent in each behavioral state per animal. Statistics, sample size and *p* values available in table.

### Noradrenergic neuromodulators regulate aggression

Persistent behavioral states are often stabilized by distinct neuromodulators. Frequently, these act antagonistically to set mutually exclusive patterns of behavior, for example by regulating the duration of opposing states like sleep and wakefulness, or hunger and satiety (*29–32*). Therefore, to explore if similar circuits are involved in establishing and maintaining the aggressive predatory states versus the non-predatory states, we screened through mutants of the four major neuromodulators using strains generated via CRISPR/Cas9 (table S1) (*33*, *34*). Mutations generated in *Ppa-tph-1, Ppa-cat-2,* and *Ppa-tbh-1* affect serotonin, dopamine, and octopamine production, respectively, while *Ppa-tdc-1* affects the production of both tyramine and octopamine, as these neuromodulators are in the same biochemical synthesis pathway (Fig. 3A). All mutants were crossed into the *Ppa-myo-2p*::RFP background and assessed using the high-throughput tracking and machine-learning pipeline previously established (Fig. 1).

**Fig. 3.**
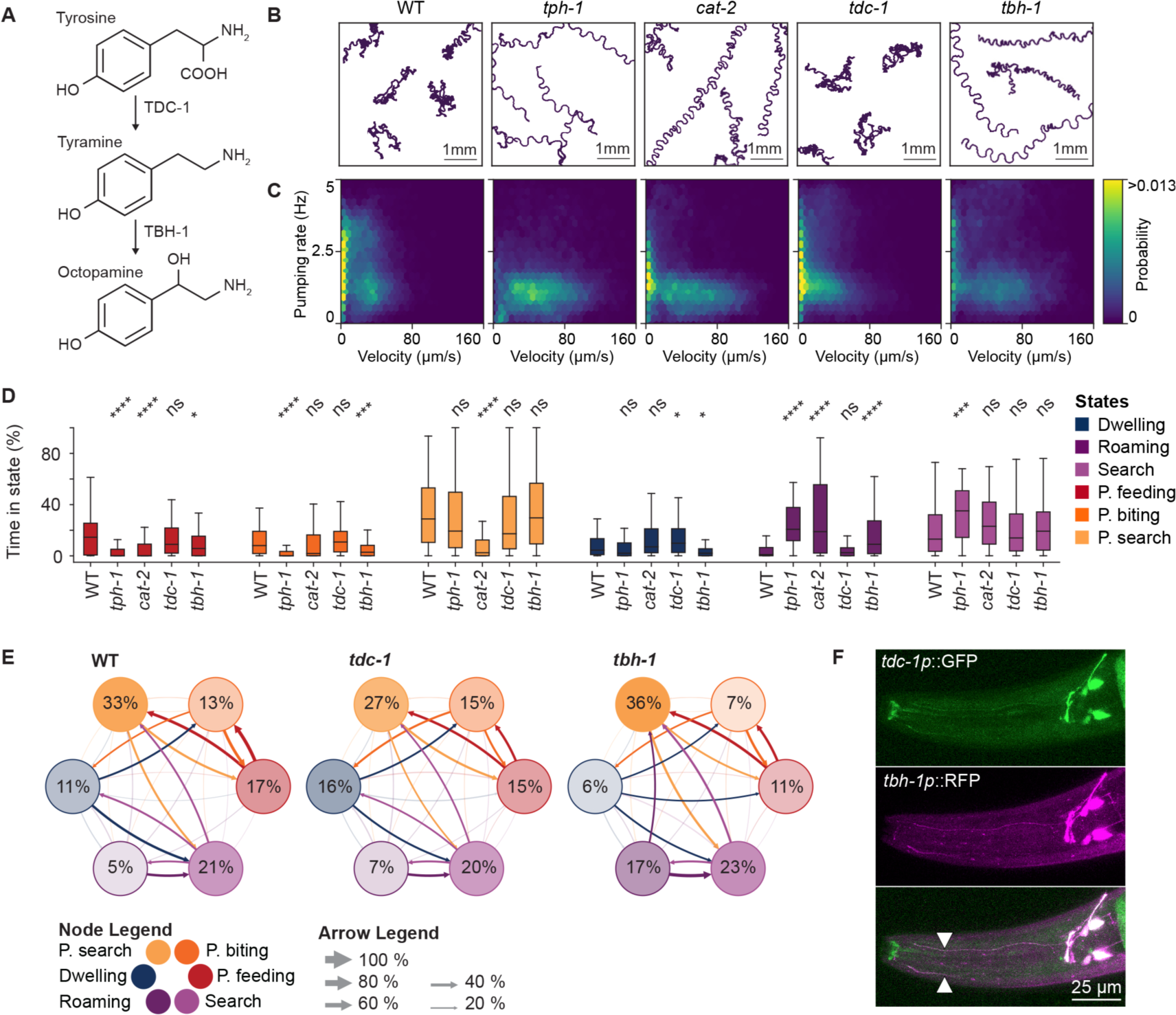
Expansion and morphological changes in noradrenergic neurons underlie predation states. (**A**) Synthesis pathway of tyramine and octopamine from the precursor tyrosine. The enzymes involved in tyramine (TDC-1) and octopamine synthesis (TBH-1) act in the same pathway. (**B**) Example animal tracks for wildtype (WT), *tph-1, cat-2, tdc-1,* and *tbh-1* mutants. (**C**) Probability density map of velocity and pumping rate for animals corresponding to the genotypes in (B). (**D**) Time spent in each behavioral state normalized to the total track duration. Significance was assessed using a Mann-Whitney U-test with a Bonferroni correction for multiple comparisons if required. (**E**) Average transition rates between behavioral states for WT, *tdc-1,* and *tbh-1* mutants on larval prey. The number in circles indicates the average state duration as in (D) and the arrow size indicates the transition rate normalized to outgoing transitions. (**F**) Expression pattern of *tbh-1p*::GFP (green) and *tdc-1p*::RFP (magenta) and colocalization. White arrows indicate neurite projecting towards the mouth from the anterior neuron pair.

In *C. elegans,* serotonin plays a multifaceted role in its behavior including regulating feeding, locomotion, foraging, egg laying, stress response, and learning and memory (*35–37*). In *P. pacificus*, previous work identified an additional predatory-specific role for this neurotransmitter in synchronizing the action of the tooth and pharyngeal pumping, proving essential for efficient predation (*34*).

Correspondingly, *Ppa-tph-1* mutants show increased roaming behaviors similar to observations in *C. elegans* and a strong decrease in predation consistent with previous findings (Fig. 3B to D and fig. S5A). Similar to *Ppa-tph-1*, the *Ppa-cat-2* dopamine-synthesis deficient animals also show a change in body posture and movement speed (Fig. 3B and C and fig. S5A) however, they do not exhibit the decrease in ‘predatory biting’ observed in *Ppa-tph-1* serotonin mutants. Instead, we see a reduction in the ‘predatory search’ and ‘predatory feeding’ states (Fig. 3D). In *C. elegans* dopamine is critical for efficient foraging when food is present (*35*, *38*), and we predict that in *P. pacificus*, dopamine may be required for initiating feeding behavior post successful kill which may relate to a food reward signal.

While the effects of serotonin and dopamine are well described in *C. elegans*, much less is known regarding the function of the noradrenergic neuromodulators’ tyramine and octopamine. Tyramine has been implicated in modulating a rapid “flight” escape response by linking head movements with locomotion and also plays a role in other long-term stress responses (*39–42*). In contrast, octopamine initiates a fasting signal facilitating exploration and optimizing foraging strategies during nutrient scarcity (*43–45*). In *P. pacificus,* we detect significantly reduced levels of the ‘predatory biting’ state in *Ppa-tbh-1* mutants and fewer transitions into this state indicating octopamine promotes aggression in *P. pacificus* (Fig. 3B to E). Strikingly, while *Ppa-tdc-1* is required for the biosynthesis of both tyramine and octopamine, in *Ppa-tdc-1* mutants predatory-associated states are maintained at wildtype levels. This indicates the additional loss of tyramine suppresses the octopamine-induced predation defect (Fig. 3B to E). A similar finding is observed in *Ppa-tdc-1; Ppa-tbh-1* double mutants which phenocopy the *Ppa-tdc-1* mutants (fig. S5B to E). In line with these data, applying exogenous tyramine to wildtype animals also induces non-predatory bouts while the addition of exogenous octopamine maintains high levels of predation (fig. S6). Thus, in *P. pacificus* there is an evolutionary divergence in the function of these neuromodulators from those previously reported in *C. elegans*. Octopamine enables robust and prolonged predatory behaviors associated with an aggressive drive while tyramine acts antagonistically to establish the docile, non-predatory state.

### Noradrenergic neurons evolved predatory adaptations

Having established the divergence in function associated with these neuromodulators in *P. pacificus,* we next explored the neural circuits associated with their biosynthesis. In *C. elegans*, the interneurons RIM and RIC represent the only tyraminergic neurons, and they express *Cel-tdc-1*, while the RIC neurons additionally express *Cel-tbh-1* and are therefore also octopaminergic (*40*). To investigate if the expression of these enzymes and the potential circuits involved in *P. pacificus* are conserved, we generated a transgenic line carrying *Ppa-tdc-1p*::GFP and *Ppa-tbh-1p*::RFP transcriptional reporters. Here, unlike in *C. elegans,* we observed two pairs of neurons expressing both *Ppa-tdc-1p*::GFP and *Ppa-tbh-1p*::RFP indicating an additional octopaminergic neuron pair in *P. pacificus*. Due to soma position and neurotransmitter expression, we putatively identify these neuron pairs as the *P. pacificus* equivalent of RIC (posterior) and RIM (anterior). However, the most anterior neuron pair is also morphologically distinct from the *C. elegans* RIM neurons with a neurite extending anteriorly towards the mouth (Fig. 3F). We hypothesize that this additional neurite represents a predatory-specific evolutionary adaptation which allows non-synaptic secretions of octopamine and tyramine along the length of the *P. pacificus* pharynx and in the vicinity of the teeth-like predatory denticles used for the killing attacks of *P. pacificus*. Therefore, in *P. pacificus* we observe two pairs of neurons expressing both neurotransmitters and with features associated with the regulation of aggressive behaviors.

### Receptor circuits are rewired across nematode evolution

Next, we attempted to elucidate the molecular mechanisms involved in generating the aggressive predatory and non-predatory states. In *C. elegans,* three octopamine receptors have been identified, *Cel-ser-3, Cel-ser-6,* and *Cel-octr-1* (*46–48*). We identified 1:1 orthologues in *P. pacificus* for all three receptors and generated CRISPR/Cas9 mutants in these genes in the *Ppa-myo-2p*::RFP strain (fig. S7, table S1 and table S2). We then assessed the predatory drives of these animals using our behavioral state model pipeline. While mutations in *Ppa-octr-1* maintained wildtype predatory biting, mutations in both *Ppa-ser-3* and *Ppa-ser-6* phenocopied the ‘predatory biting’ state observed in *Ppa-tbh-1* mutant (Fig. 4A to C and fig. S8A). Therefore, both *Ppa-ser-3* and *Ppa-ser-6* are required to establish efficient *P. pacificus* predatory bouts through octopamine signaling. Similarly, there are four known tyramine receptors in *C. elegans, Cel-tyra-2, Cel-tyra-3, Cel-ser-2,* and *Cel-lgc-55* (*47*, *49–51*). In order to identify potential tyramine receptors involved in this pathway, we also identified 1:1 orthologues in *P. pacificus* for all four of these receptors and generated corresponding CRISPR/Cas9 mutants in the *Ppa-tbh-1; Ppa-myo-2p*::RFP strain to determine if any rescued the *Ppa-tbh-1* reduced-killing phenotype (fig. S7, table S1 and table S2). Mutations in *Ppa-tyra-2, Ppa-tyra-3,* and *Ppa-ser-2* as well as the corresponding triple mutant maintained low levels of predatory aggression similar to the *Ppa-tbh-1.* However, in *Ppa-lgc-55* mutants, predation was restored to similar levels as seen in wildtype and *Ppa-tdc-1* mutants (Fig. 4 D to F and fig. S8, B to E). Additionally, we observe a substantial increase in the ‘predatory search’ behavior in *Ppa-tyra-3* mutants indicating a potential role for *Ppa-tyra-3* in regulating this behavioral state (Fig. 4 D to F and fig. S8 B). Thus, two octopamine receptors and a single tyramine receptor are required to mediate the predatory states and associated aggressive transitions.

**Fig. 4.**
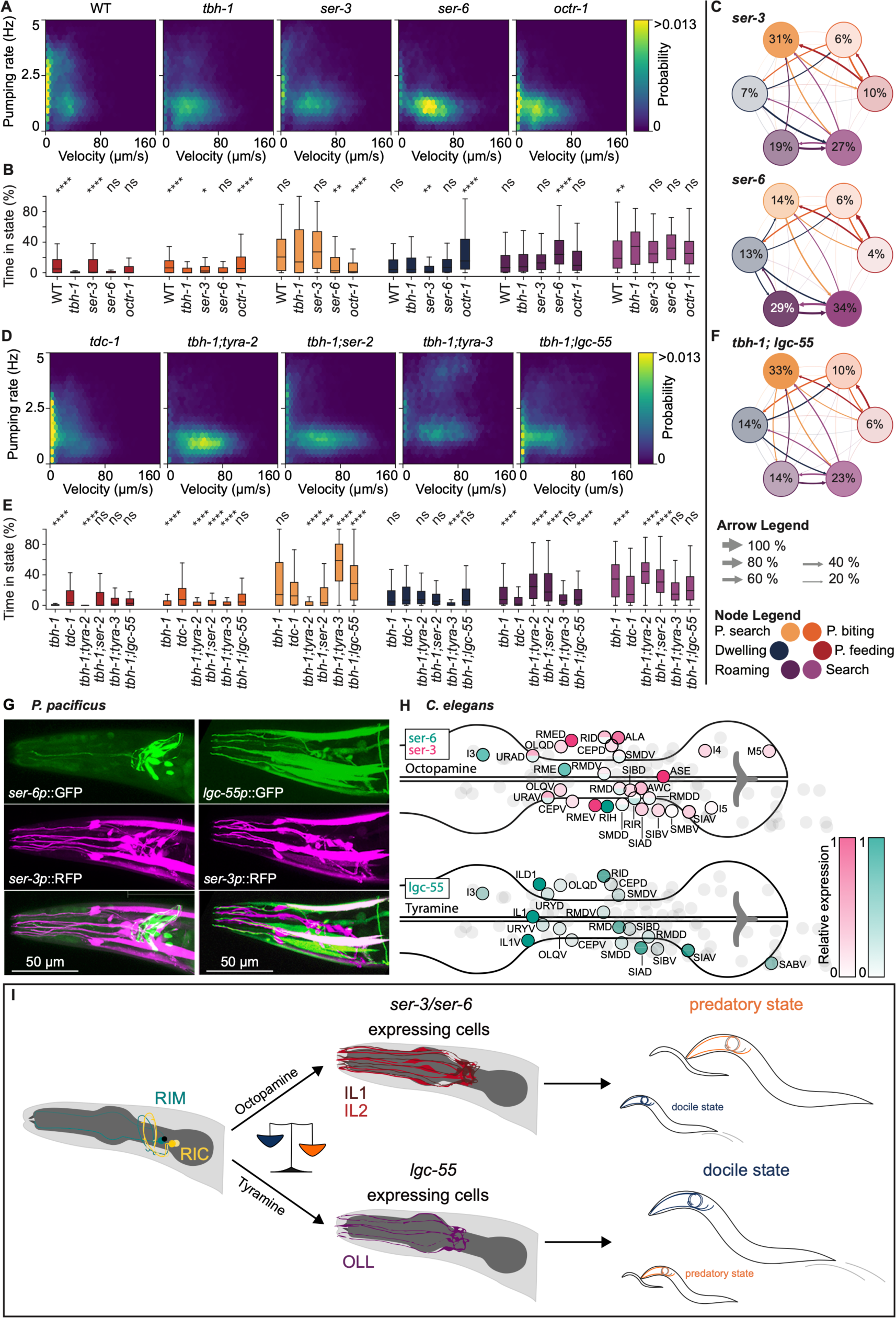
Octopamine and tyramine receptors in sensory neurons gate aggressive state entry and exit. (**A**) Probability density map of velocity and pumping rate for WT, *tbh-1* and the octopamine receptors *ser-3*, *ser-6* and *octr-1*. *ser-3* and *ser-6* phenocopy *tbh-1*. (**B**) Relative time in each behavioral state for all genotypes in (A). (**C**) Average transition rates between behavioral states for *ser-3* and *ser-6*. The number in circles indicates the average state duration as in (B), and the arrow size indicates the transition rate normalized to outgoing transitions. (**D**) Probability density map of velocity and pumping rate for *tdc-1* and the tyramine receptors *tyra-2, ser-2, tyra-3*, and *lgc-55*, all in the *tbh-1* background. (**E**) Relative time in each behavioral state for all genotypes in (D). (**F**) Average transition rates between behavioral states for *tbh-1; lgc-55.* The number in circles indicates the average state duration as in (E) and the arrow size indicates the transition rate normalized to outgoing transitions. (**G**) Expression pattern for the octopamine receptors *ser-3* (magenta) and *ser-6* (green), as well as the tyramine receptor *lgc-55* (green). (**H**) Expression pattern of the *C. elegans* receptors *ser-3* (magenta) and *ser-6* (green), and *lgc-55* (green) in the animal head, based on data from the CeNGEN single-cell transcriptome analysis. Gray dots represent neurons that don’t show expression. (**I**) Proposed model of the octopamine/tyramine gating of the sensory neurons that drives aggressive versus docile behavioral states.

Finally, to identify the neurons governing behavioral state switching we generated reporter lines for all three of the receptors associated with the ‘predatory biting’ states. The octopamine receptor *Ppa-ser-3* is expressed in the neck muscles, several head neurons, and strikingly the IL2 and IL1 head sensory neurons with their distinctive neurites projecting to the worm’s nose (*52*). *Ppa-ser-6* is expressed in a non-overlapping set of head neurons which also includes a neuron pair with anterior processes (Fig. 4G).

A strain expressing *Ppa-tbh-1p*::RFP and *Ppa-ser-6p*::GFP revealed co-expression of this receptor in the neuron pair we previously putatively identified as RIM (Fig. 3F and fig. S9A). This may indicate a potential feedback mechanism acting through these neurons regulating the release of octopamine and tyramine around the pharynx and teeth-like structures. As the *Cel-ser-6* receptor couples to the excitatory G_alphaS pathway, this suggests a positive auto-receptor motif where octopamine release increases neuronal excitation and stimulates further octopamine release along the pharynx and teeth-like structures (*48*). Such a motif can induce bistable state-switches, which may underlie the observation of persistent and robust predatory episodes. For the tyraminergic receptor *Ppa-lgc-55,* we again observe robust expression in the neck muscles which is similar to observations in *C. elegans* (*48*), but we also detect strong expression in a distinct set of head sensory neurons separate from the octopamine receptor expressing IL2s and IL1s (Fig. 4G). These are putatively identified as the OL neurons based on neurite morphology and soma placement (*51*). In contrast, using the CeNGEN dataset (*53*), we found these three receptors in *C. elegans* are expressed throughout a large subset of head neurons including sensory, inter, and motor neurons. However, there is minimal overlap with any of the head sensory neurons we observe in *P. pacificus* (Fig. 4H). Therefore, in *P. pacificus* octopamine and tyramine receptors associated with aggressive behavioral states are localized to unique head sensory neurons many of which project sensory endings into the environment (fig. S9B). Moreover, these sensory endings represent the first point of contact between predator and prey and as such are prime candidates for prey detection neurons. Taken together, we envisage a model whereby octopamine and tyramine antagonistically regulate aggressive state-switching (Fig 4I). This occurs through state-dependent gating of head sensory neurons which, when stimulated, can be utilized to detect the presence of a contact event with prey and result in an aggressive predatory attack.

## Discussion

Reconstructing the evolutionary changes generating novel behaviors is dependent on combining neurobiological and behavioral studies between phylogenetically characterized species. Accordingly, the divergent behaviors observed between *C. elegans* and *P. pacificus* are well suited to uncovering innovations associated with behavioral evolution. In our study, we focused on the evolution of *P. pacificus* aggression. In *P. pacificus,* aggression is a state associated with the regulation of multiple behaviors including predation, cannibalism, and territoriality (*11–14*). We found that *P. pacificus* aggression is regulated through the actions of octopamine and tyramine, which share many similarities with known attributes of aggression in several other invertebrate species. This includes territorial behaviors and nestmate recognition in social insects, intraspecific and intersexual cannibalism in arachnids, and fighting in crustaceans (*54*). Similarly, in *D. melanogaster* aggression has been shown to also depend on a complex system of neurotransmitters including octopamine as well as other hormones and neuropeptides (*55–62*). In *C. elegans* no aggression has been reported, and instead octopamine and tyramine appear to have evolved independent functions (*40*) with tyramine broadly regulating the escape response and octopamine inducing fasting-associated behaviors (*39*, *41–44*). In contrast, our study shows that during the evolution of *P. pacificus,* the function of these neuromodulators has become highly interlinked and they act antagonistically to instead regulate its predatory aggression.

Coinciding with this, we also find diversity in the neuronal wiring between *C. elegans* and *P. pacificus* associated with the evolution of *P. pacificus* aggression. Sensory neurons are often thought to represent the regions of greatest plasticity and are most frequently associated with behavioral changes in several other species (*63–66*). However, in our study we find evolutionary changes are evident across the entire neural circuit associated with *P. pacificus* aggression. Specifically, we find an additional pair of octopaminergic neurons which also possess key morphological differences including neurites capable of secreting these neuromodulators along the length of the pharynx. This likely represents a predation-specific adaptation to deliver neuromodulators closer to their site of action. Additionally, the distribution of octopamine and tyramine receptors has greatly diverged across the evolution of these two species with *P. pacificus* acquiring distinct receptor expression throughout many of its head sensory neurons. These neurons are strong candidates for prey sensing due to their exposure to the external environment. Accordingly, we predict the *P. pacificus* noradrenergic system may regulate the activity of these neurons and facilitate the detection of prey through as yet unknown mechanosensory and / or chemosensory receptors found in its environmentally exposed sensory organs. Thus, noradrenergic circuits balance aggressive behavioral states in nematodes and are associated with the evolution of complex behavioral traits.

## Materials and Methods

### Animal handling and maintenance

The strains used in this study are shown in table S1. *C. elegans* and *P. pacificus* were maintained at 20 °C on Nematode Growth Medium (NGM) agar plates containing *Escherichia coli* OP50.

### Transgenic animals

To generate the *Ppa-myo-2p*::RFP (JWL27) strain, we used the previously established protocol (*67*). This construct was generated by PCR amplification of a 1231 bp upstream region in front of the first predicted ATG start codon of *Ppa-myo-2* and subsequent cloning into the pZH009 containing the codon-optimized red fluorescent protein (TurboRFP) plasmid. NEBuilder HiFi DNA Assembly Master Mix (New England Biolabs) was employed to perform cloning. To generate the transcriptional reporters of *Ppa-tbh-1*, *Ppa-tdc-1, Ppa-ser-3, Ppa-ser-6,* and *Ppa-lgc-55*, we cloned the upstream regions before their predicted start codon including: *Ppa-tbh-1*: 1958 bp; *Ppa-tdc-1*: 1585 bp, *Ppa-ser-3*: 1996 bp, *Ppa-ser-6*: 1996 bp and *Ppa-lgc-55*: 1917 bp to drive expression of the codon-optimized TurboRFP or GFP as required. Each injection mix contained 10 ng/µl of the PstI-HF-digested reporter plasmids, 10 ng/µl of the PstI-HF-digested *Ppa-egl-20p*::GFP plasmid as co-injection marker, and 60 ng/µl of the PstI-HF-digested genomic carrier DNA. The mix was injected in the gonads of young adults. The transgenic animals were screened using an epi-fluorescence microscope (Axio Zoom V16; Zeiss). The fluorescent images of the transgenic animals were obtained using a Leica SP8 confocal microscope.

### Transgenic line integration

*Ppa-myo-2p*::RFP was integrated into the *P. pacificus* genome as previously described (*68*). Briefly, 10 NGM plates each containing 20 fluorescent *Ppa-myo-2p*::RFP animals were exposed to UV irradiation at 0.050 J/cm^2^ using a UV Crosslinker (CL-3000 Analytik Jena). After 3-4 days F1 fluorescent animals were singled out onto 120 individual culture plates and after another 3-4 days the F2 progeny screened for possible integration events. This was detected by observing an increase in the number of fluorescent animals to ≥75% of the population. Individual animals from these plates were isolated and screened for consistent 100% transmission. Integrated lines were subsequently outcrossed 4x to remove potential mutations caused by UV exposure.

### Behavioral imaging

*Ppa-myo-2p*::RFP animals were recorded at 1x effective magnification using an epi-fluorescence microscope (Axio Zoom V16; Zeiss) as described in (*18*). Recordings were made via a Basler camera (acA3088-57um; BASLER) with 15 ms exposure time. Animals were imaged at 30 fps for 10 min unless otherwise indicated. All animals that were in the field-of-view for at least 60 seconds were included in the analysis.

For tracking of animals on predatory assays, *C. elegans* prey were firstly maintained on bacteria OP50 until freshly starved, resulting in an abundance of young larvae. These plates were washed with M9, passed through two 20 µm filters, centrifuged and deposited onto the assay plate by pipetting 4 µl of worm pellet onto a 6 cm NGM unseeded plate. A copper arena (1.5 cm x 1.5 cm) was placed in the middle of the assay plate to constrain predators in the recording field. 40 young adult *P. pacificus* predators (Eurystomatous mouth form) were starved for 2 hours and then added to assay plates inside the arena. After a recovery period of 15 mins on the assay plate, their behaviors were recorded for 10 mins.

For tracking of animals on bacterial assays, 300 µl of *E. coli* OP50 culture was spotted onto an empty 6 cm NGM plate 24 h before the assay. A copper arena (1.5cm x 1.5cm) was placed in the middle of the assay plate to contain the *P. pacificus* in the recording field. 40 young adult predators were starved for 2 h and then added to assay plates inside the arena. After a recovery period of 15 mins on the assay plate their behaviors were recorded for 10 mins.

### Automated behavioral tracking

Animals were tracked using the custom python analysis package *PharaGlow* (*18*). PharaGlow performs a three-step analysis: 1. center of mass tracking and collision detection, 2. linking detected objects to trajectories, and 3. extracting centerline, contour, width, and other parameters of the shape to allow extracting pharyngeal pumping events. The generated files contain the position, and the straightened images which are further processed to extract the behavioral measures. Post-processing was applied according to Bonnard et al., 2022 (*18*) with minor modifications. To obtain pumping traces from straightened animals, the inverted skew of intensity is calculated for each frame per animal (Fig. 1 C). This metric is sensitive to the opening of the pharyngeal lumen and pharyngeal contractions. Peaks in the resulting trace correspond to pumping events.

### Feature Engineering

Videos of animals with labeled pharyngeal muscles were processed using PharaGlow (*18*), which provided a set of initial behavioral features (center of mass coordinates (CMS), the centerline, the pumping rate, the skew of the fluorescence intensity distribution, which relates to pumping contractions. From these basic features, we calculated two additional derived features. Using the CMS coordinates (x(t), y(t)) velocity was calculated with a time shift of dt = 60 frames (= 2 s). To obtain a description of head motion, we calculated the angle between the movement direction of the CMS and nose tip as θ = arccos (*v_CL_v_CMS_*), where *v_CL_* is the unit vector of the centerline between the 1^st^ and 5^th^ coordinate along the 100 equidistantly sampled points of the centerline and *v_CMS_* is the unit vector of the center of mass coordinates between two consecutive frames. To capture frequency changes in these features, we apply a wavelet transform using pywt with a gaus5 wavelet and extract the maximum frequency for each feature over time. The head angle and the skew which relates to the faster pumping motion were transformed with the following range of pseudo-frequencies (0.3-5.0 Hz). From the wavelet transformations the maximum frequency was extracted and included as a feature. Only for skew two wavelet transforms from scale 11.2 and 3.8 were directly included additionally, which translates to a pseudo-frequency of ∼1.3 and ∼3.9 Hz, frequencies relevant for feeding behavior. For velocity only one wavelet transform with the scale 18.7 was included, which translates to ∼0.8 Hz. This transformation resulted in a total set of 9 features.

### Data curation

During recordings with many animals, a small subset of frames includes overlapping animals. We detect and remove these instances in the data by removing frames with an area higher than 1.5 times the mean area within one recording.

### Manual annotation

An expert human annotator generated labels for a subset of frames following the naming conventions for behavior in Wilecki et al., (*16*) and Okumura et al., (*34*). Using LabelStudio pumping rate and velocity data was displayed and the annotator marked sequences corresponding to ‘biting’, ‘feeding’, ‘exploration’ or ‘quiescence’. The labels were verified by playing the corresponding movie of the animal. For the unlabeled recordings, we automatically set labels encoding the recording condition, namely “on OP50” or “on *C. elegans* larvae”.

### Preprocessing

The original features were preprocessed and downsampled before analysis using the following pipeline implemented in sklearn. First, additional lagged features were created from the base features (velocity, instantaneous pumping rate, head angle, mean pumping rate) by shifting the individual dataset by 5, 10 and 15 frames (with 30 fps) forward or backward in time. This allowed us to include a short history in an otherwise memory-less pipeline. Subsequently, features were averaged using a rolling window of 30 frames (1 s) and downsampled to 1 frame per second. Labels were downsampled by using the modal value within 30 frames (1 s). The first and last second of each recording were truncated, in order to eliminate effects of smoothing artifacts. The data was then transformed with a Yeo-Johnson power transform to improve normality and normalized with robust scaling. The entire pipeline was fitted on the training dataset and also applied during prediction of novel data.

### Dimensionality Reduction and Clustering

The preprocessed training data with N = 106 animals from WT recordings on larvae and bacteria was embedded in three dimensions using UMAP (umap module, Python) using the parameters: n_neighbors = 70, min_dist = 0. repulsion_strength = 4, negative_sample_rate = 15, disconnection_distance = 0.85. n_components = 3. The resulting embedding was clustered with HDBSCAN. The number of clusters was determined using a silhouette score. To assign human-interpretable behavioral states to the clusters, the overlap between clusters and manual-labels was calculated (Fig. S2C) and cluster labels were verified by inspecting the feature distributions within the different clusters. Finally, cluster labels were spot-checked using the video data.

### Behavioral state classification

Behavioral feature data with cluster labels found by embedding and clustering were used to train an XGBoost Classifier on the preprocessed data. Nine of the 106 videos were held-out as test set and were not used for training the classifier. These videos were selected for their similarity in label distribution compared to the training set. Bayesian optimization was used to find optimal hyperparameters for the classifier using cross validation. Training data was split using a stratified group shuffle split, whereby each recording was constituting a group. After the best hyperparameters were found, a model was trained on the complete training data using the optimal hyperparameters. Next, the performance of the model was evaluated with the prediction of the test set in comparison with the cluster labels. For this, standard model performance metrics were calculated (Fig. 1, and fig. S3A). Before prediction the steps feature engineering and preprocessing are applied to the novel data. To ensure reliability of the predictions, predictions with a probability of less than 50% are set to “None”.

### Statistical analysis

For the statistical analysis of the state predictions (relative time in state, mean bout duration, transition rates), the two-tailed Mann-Whitney-U test was used. The statistical analysis was performed on each condition and state to its respective control. A Bonferroni correction was applied to take into account that the probability of a false positive significant test rises with the number of compared conditions. Thus, the *p* values reported here are corrected for the number of comparisons made to the respective control. All raw *p* values and sample sizes are available in table S3. In all figures * denotes a significance value between 0.05 and 0.01, ** a significance value between 0.01 and 0.001, *** a significance value between 0.001 and 0.0001, and **** a significance below 0.0001.

### Generation of CRISPR/Cas9 induced mutations

The procedure for CRISPR/Cas9 mutagenesis was based on previously described *P. pacificus* methods (*69*). Briefly, gene-specific CRISPR RNAs (crRNAs) were designed to target early predicted exons in all target genes and synthesized by Integrated DNA Technologies (IDT). These were then fused to tracrRNA (IDT) at 95°C for 5 minutes before cooling to room temperature and annealing. The hybridization result was coupled with purified Cas9 protein (IDT). After 5 minutes of incubation at room temperature, TE buffer was added to achieve a final concentration of 18.1 μM for sgRNA and 12.5 μM for Cas9. The mix was injected in the gonads of young adults. P0 worms were discarded 12–24 h after microinjection. After two days, 96 F1 progeny from P0 plates were singled out and allowed to lay eggs for 24 hours. The genotype of the F1 animals were subsequently analyzed via Sanger sequencing. For detailed information about the mutants created during this study see fig S7 and table S1. All sgRNAs and the primers used in this study to generate mutants can be found in table S2.

### Pharmacological experiment

To quantify the effects of exogenous octopamine and tyramine, predatory feeding assays were performed on agar plates which are supplemented with 2 mM octopamine and tyramine respectively. Tyramine and octopamine plates were prepared by adding tyramine-HCl (Sigma) and octopamine-HCl (Sigma) to a concentration of 2 mM to freshly autoclaved NGM that was cooled down to 55 ℃ prior to use. *Ppa-myo-2p*::RFP animals were picked onto either unseeded octopamine and tyramine containing plates and starved for two h. These animals were subsequently, added to standard assay plates with *C. elegans* larvae contained in a copper arena and also containing octopamine or tyramine.

### Egg-laying assay

Three-day-old adult worms were placed on individual plates seeded with OP50. For the following day, the number of eggs was manually counted and the worms were transferred onto new plates. The eggs were counted for 5 days per individual worm after which only unfertilized eggs were laid.

### Manual predation assays

Assays were conducted as previously reported in Wilecki and Lightfoot et al., 2015 (*16*). Briefly, Prey were maintained on NGM plates seeded with OP50 bacteria until freshly starved, resulting in an abundance of young larvae. These plates were washed with M9 and passed through two 20 μm filters to isolate larvae. 1.0 μl of *C*. *elegans* larval pellet was transferred onto a 6 cm NGM unseeded plate. Five predatory nematodes were screened for the appropriate mouth morph and added to assay plates for prey assays. After 2 h the plate was screened and corpses were counted manually.

### Worm size measurement

*S*ynchronized J2 larvae were placed into NGM plates with bacteria. They are transferred from assay plates to NGM plates without bacteria during different developmental points (24, 48 and 72 h). Bright field images of the worms were taken using an epi-fluorescence microscope (Axio Zoom V16; Zeiss) and the Basler camera (acA3088-57um; BASLER). Images were analyzed using the Wormsizer plugin for Image J/Fiji. Wormsizer detects and measures the area of the worms.

### Mouth-form phenotyping

Mouth-form phenotyping was performed as previously reported in Ragsdale et al., 2013 (*15*). In brief, NGM plates with synchronized young adults were placed onto a stereomicroscope with high magnification (×150). 100 worms were screened for the mouth form per strain. The eurystomatous (Eu) mouth form was determined by the presence of a wide mouth, whereas the stenostomatous (St) forms were determined by a narrow mouth. Eu young adult worms were picked for predation assays.

### Ppa-myo2p::RFP copy number quantification

Thirty J4 / young adult hermaphrodites from a non-starved plate were picked for DNA extraction using a NEB Monarch DNA kit (final elution volume 35 µl). A whole plate of non-starved worms was used for RNA extraction using a Zymo RNA mini kit (final elution volume 30 µl).

To determine the copy number of *Ppa-myo-2p*::RFP in the UV integrated transgenic line (JWL27) and relative expression of this construct, qPCR and RT-qPCR assays were conducted. For copy number, primers were designed to amplify a part of RFP sequence and compared to two known single copy gene sequences *Ppa-gpd-3* and *Ppa-csg-1*. As *Ppa-myo-2p*::RFP is a transcriptional reporter no part of the *Ppa-myo-2* gene sequence is included in the reporter line. Therefore, to determine the relative expression *Ppa-myo-2p::*RFP, the expression of RFP was compared to that of the native *Ppa-myo-2* exon gene sequence. qPCR primers specific for RFP were Forward GGAGAGGGAAAGCCTTACGAGG and Reverse GAATCCCTCAGGGAAAGACTGC. Two pairs of gene specific *Ppa-myo-2* primer were used. These were pair 1: forward CGAAGAAGAACGTGTGGGTG and reverse TACCTCATTGCCGGGACCTC. Pair 2: forward AGGAGACAAAGGGAGACACG and reverse GGGTTCATCTCCTGCACTTGG.

For quantification, DNA samples were 1:10 diluted with water, while RNA samples were not diluted. Three technical replicates and two biological replicates were conducted.

## Acknowledgments

We would like to thank Ralf Sommer for discussions and critical reading of the manuscript, Wolfgang Bönigk for transgenic cloning assistance (Genetics Facility, MPI for Neurobiology of Behavior-caesar), and Jürgen Berger for SEM imaging (MPI for Biology). Additionally, we wish to thank the Sommer lab for *P. pacificus* wildtype strain PS312 (MPI for Biology, Tübingen), Finally, the *C. elegans* wildtype N2 strain was provided by the CGC, which is funded by NIH Office of Research Infrastructure Programs (P40 OD010440).

## Author contributions

Conceptualization: GGE, LB, MS, JWL

Methodology: GGE, LB, MS, JWL

Investigation: GGE, LB, MR, FH, JL, LA, DLG,NZ, ZH, MO, MS, JWL

Visualization: GGE, LB, MS

Funding acquisition: MS, JWL

Project administration: MS, JWL

Supervision: MS, JWL

Writing – original draft: MS, JWL

Writing – review & editing: DLG, MS, JWL

## Data and materials availability

All data are available in the main text or the supplementary materials.

