## Supplemental Files for "Predatory aggression evolved through adaptations to noradrenergic circuits"

Supp. 1

A

B

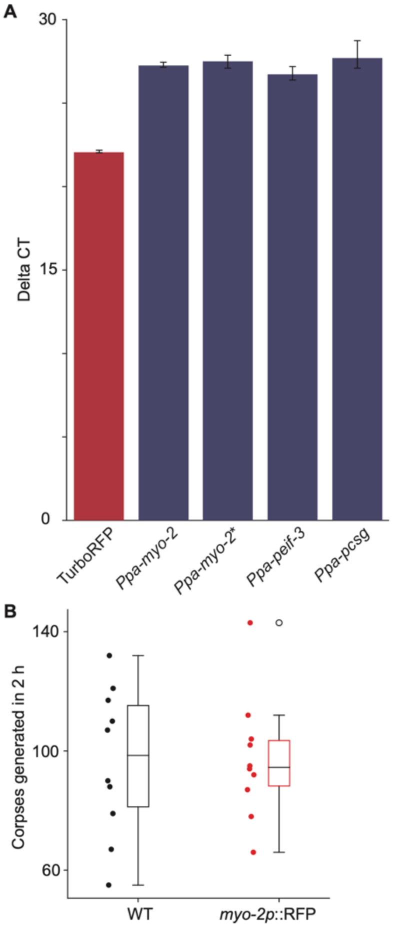


C

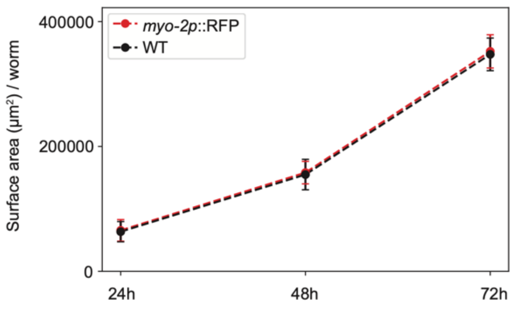


D

| | WT | *myo-2*::RFP |
|---|---|---|
| Proportion of Eu animals | 96% | 98% |

E

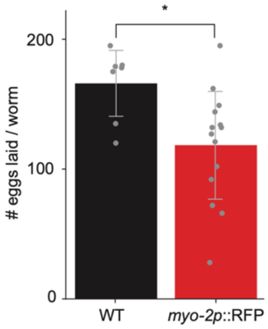

**Fig. S1. Integration of *Ppa-myo-2p*::RFP**

(**A**) qPCR analysis of RFP copy number. RFP qPCR reveals 5.5 less cycles of RFP compared to two different primer pairs specific to the *Ppa-myo-2* gene and two known single copy genes *Ppa-csg-1* and *Ppa-eif-3* indicating ~32-45 copies of *Ppa-myo-2p*::RFP are integrated into the *P. pacificus* genome in strain JWL27. (**B**) Predatory behavior is unaffected in the *Ppa-myo-2p*::RFP integrated line. Standard corpse assays with 5 young adult *P. pacificus* predators placed onto assay plates containing an abundance of *C. elegans* larvae to predate on for 2 h. 10 replicates were conducted for wild type and *Ppa-myo-2p*::RFP. (**C**) Growth rate and (**D**) phenotypically plastic mouth morph frequency are unaffected in the *Ppa-myo-2p*::RFP integration line. Eurystomatous (Eu) is the predatory mouth form in *P. pacificus*. (**E**) There is a small reduction in fecundity associated with the *Ppa-myo-2p*::RFP integration. Significance was assessed using a Mann-Whitney U-test ($p = 0.015$).

Supp. 2

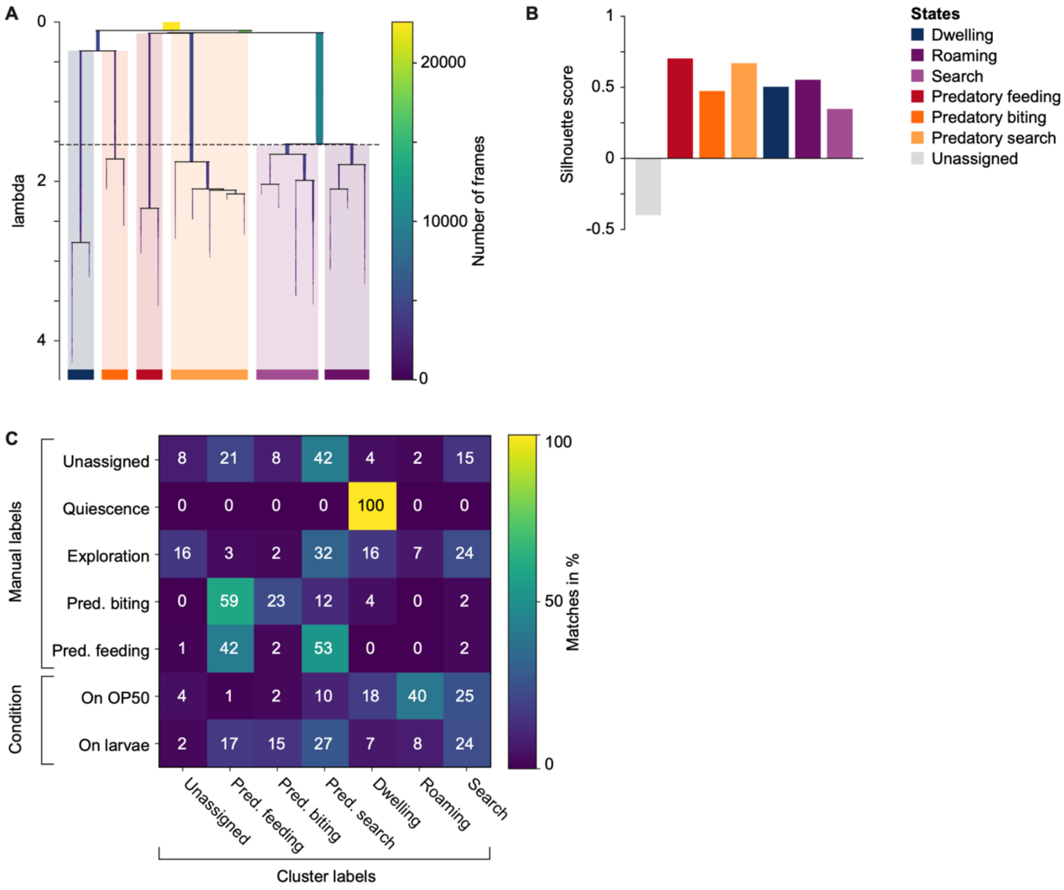


| | Unassigned | Pred. feeding | Pred. biting | Pred. search | Dwelling | Roaming | Search |
|---|---|---|---|---|---|---|---|
| Unassigned | 8 | 21 | 8 | 42 | 4 | 2 | 15 |
| Quiescence | 0 | 0 | 0 | 0 | 100 | 0 | 0 |
| Exploration | 16 | 3 | 2 | 32 | 16 | 7 | 24 |
| Pred. biting | 0 | 59 | 23 | 12 | 4 | 0 | 2 |
| Pred. feeding | 1 | 42 | 2 | 53 | 0 | 0 | 2 |
| On OP50 | 4 | 1 | 2 | 10 | 18 | 40 | 25 |
| On larvae | 2 | 17 | 15 | 27 | 7 | 8 | 24 |

**Fig. S2. Clustering of behavioral data and state identification**

(A) Hierarchical clustering of UMAP-embedded behavioral features. Colors correspond to the individual clusters that are later identified as behavioral states. The horizontal line indicates where the hierarchical tree was cut. (**B**) Silhouette scores for each cluster. (**C**) Confusion matrix between the human expert annotator, condition, and the cluster labels. Unlabeled data is included by condition as either 'On larvae' or 'On OP50'. The latter was used to identify condition-specific states.

Supp. 3

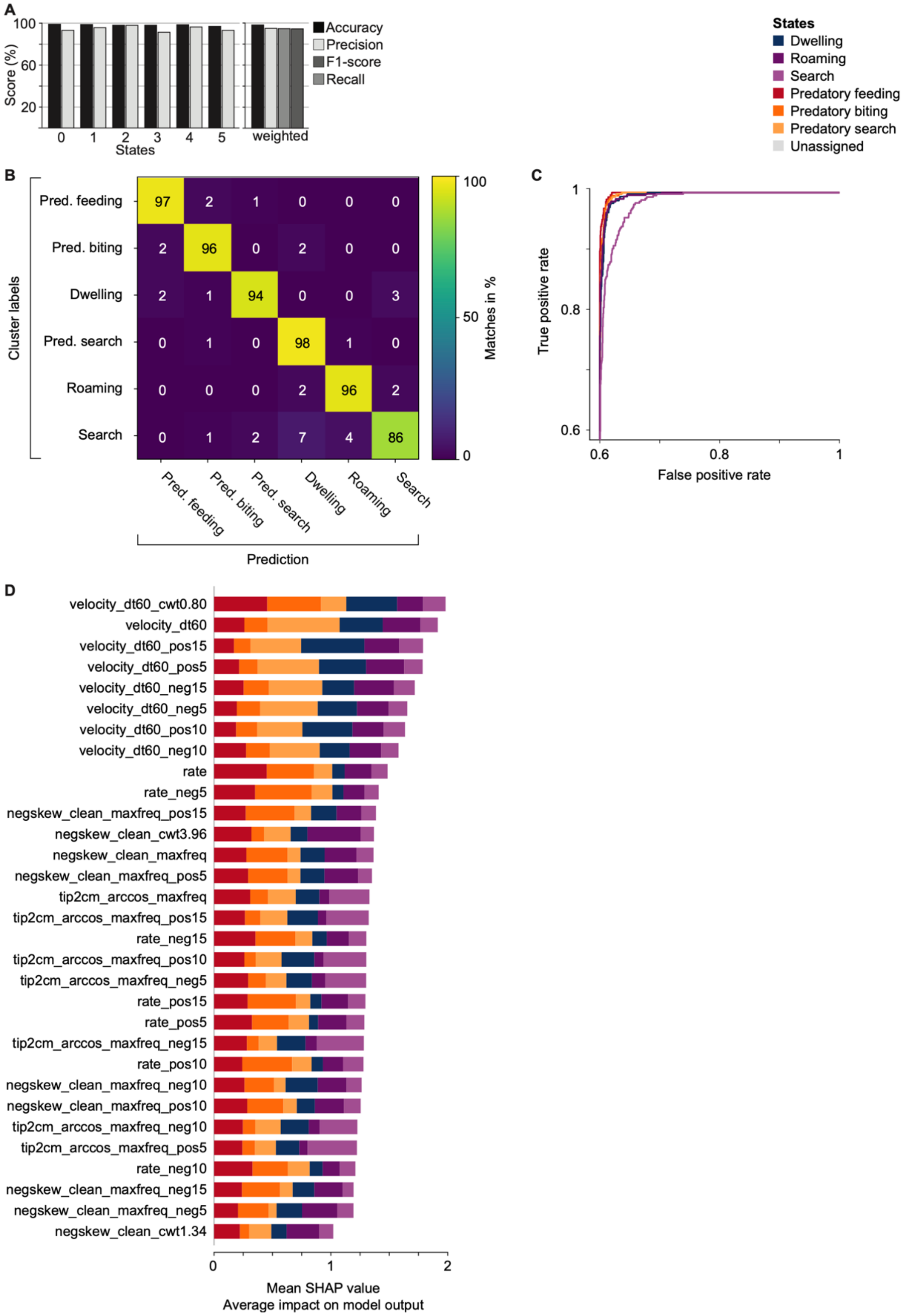

A
100
60
20
Score (%)
0
1
2
3
4
5
States
weighted
Accuracy
Precision
F1-score
Recall
States
Dwelling
Roaming
Search
Predatory feeding
Predatory biting
Predatory search
Unassigned
B
Cluster labels
Pred. feeding
Pred. biting
Dwelling
Pred. search
Roaming
Search
97 2 1 0 0 0
2 96 0 2 0 0
2 1 94 0 0 3
0 1 0 98 1 0
0 0 0 2 96 2
0 1 2 7 4 86
Pred. feeding
Pred. biting
Pred. search
Dwelling
Roaming
Search
Prediction
100
50
0
Matches in %
C
1
0.8
0.6
True positive rate
0.6
0.8
1
False positive rate
D
velocity_dt60_cwt0.80
velocity_dt60
velocity_dt60_pos15
velocity_dt60_pos5
velocity_dt60_neg15
velocity_dt60_neg5
velocity_dt60_pos10
velocity_dt60_neg10
rate
rate_neg5
negskew_clean_maxfreq_pos15
negskew_clean_cwt3.96
negskew_clean_maxfreq
negskew_clean_maxfreq_pos5
tip2cm_arccos_maxfreq
tip2cm_arccos_maxfreq_pos15
rate_neg15
tip2cm_arccos_maxfreq_pos10
tip2cm_arccos_maxfreq_neg5
rate_pos15
rate_pos5
tip2cm_arccos_maxfreq_neg15
rate_pos10
negskew_clean_maxfreq_neg10
negskew_clean_maxfreq_pos10
tip2cm_arccos_maxfreq_neg10
tip2cm_arccos_maxfreq_pos5
rate_neg10
negskew_clean_maxfreq_neg15
negskew_clean_maxfreq_neg5
negskew_clean_cwt1.34
0
1
2
Mean SHAP value
Average impact on model output

**Fig. S3. Evaluation of the XGBoost classifier model**

**(**A) Accuracy, precision, recall, and F1-score per state and the weighted average per label. (**B**) Confusion matrix comparing the cluster labels and the predicted labels using the model. (**C**) ROC curve comparing the false-positive versus false-negative rates per state. (**D**) SHAP analysis for each feature included in the machine-learning pipeline.

Supp. 4

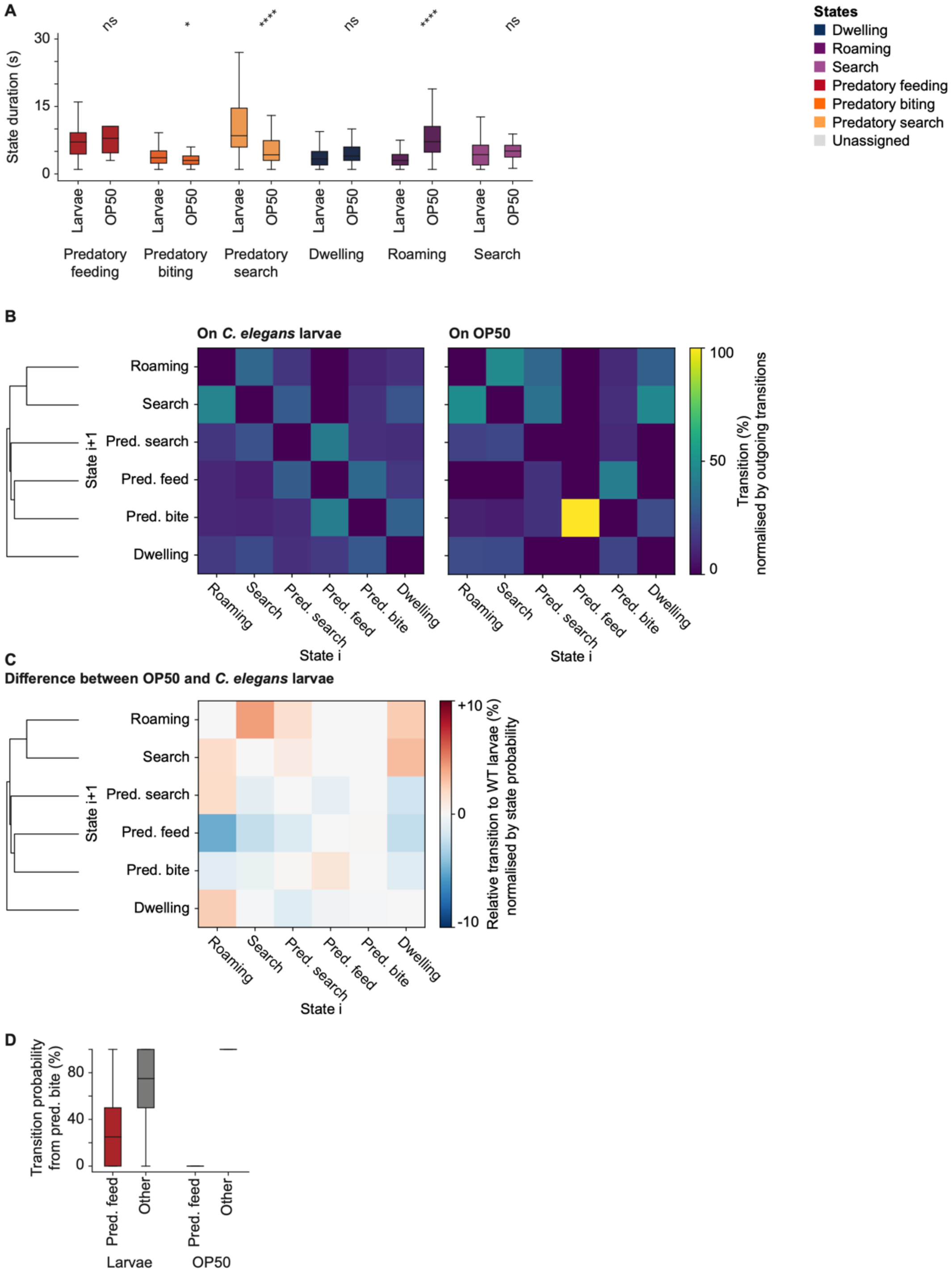

A
State duration (s)
30
15
0
ns
*
****
ns
****
ns
Larvae
OP50
Predatory feeding
Predatory biting
Predatory search
Dwelling
Roaming
Search
States
Dwelling
Roaming
Search
Predatory feeding
Predatory biting
Predatory search
Unassigned
B
On C. elegans larvae
On OP50
Roaming
Search
Pred. search
Pred. feed
Pred. bite
Dwelling
State i+1
State i
100
50
0
Transition (%) normalised by outgoing transitions
C
Difference between OP50 and C. elegans larvae
+10
0
-10
Relative transition to WT larvae (%) normalised by state probability
D
Transition probability from pred. bite (%)
80
40
0
Pred. feed
Other
Larvae
OP50

**Fig. S4. Behavioral state prediction for animals in predatory or a bacterial food context**

(**A**) Average state duration for animals on OP50 or on prey larvae. (**B**) Hierarchical clustering of the transition rates between states for each food context. (**C**) Difference in transition matrices between the different contexts, scaled by the probability of each state. Note the increase in roaming-dwelling transitions, and the search behavior on OP50, compared to a reduction of transitions into predatory behaviors like predatory search and predatory feeding. (**D**) Transition probability that a *P. pacificus* animal transitions from 'predatory biting' to 'predatory feeding' (nutritional drive) or alternatively into any other states. Analysis conducted with *P. pacificus* surrounded with potential prey or a lawn of OP50.

Supp. 5

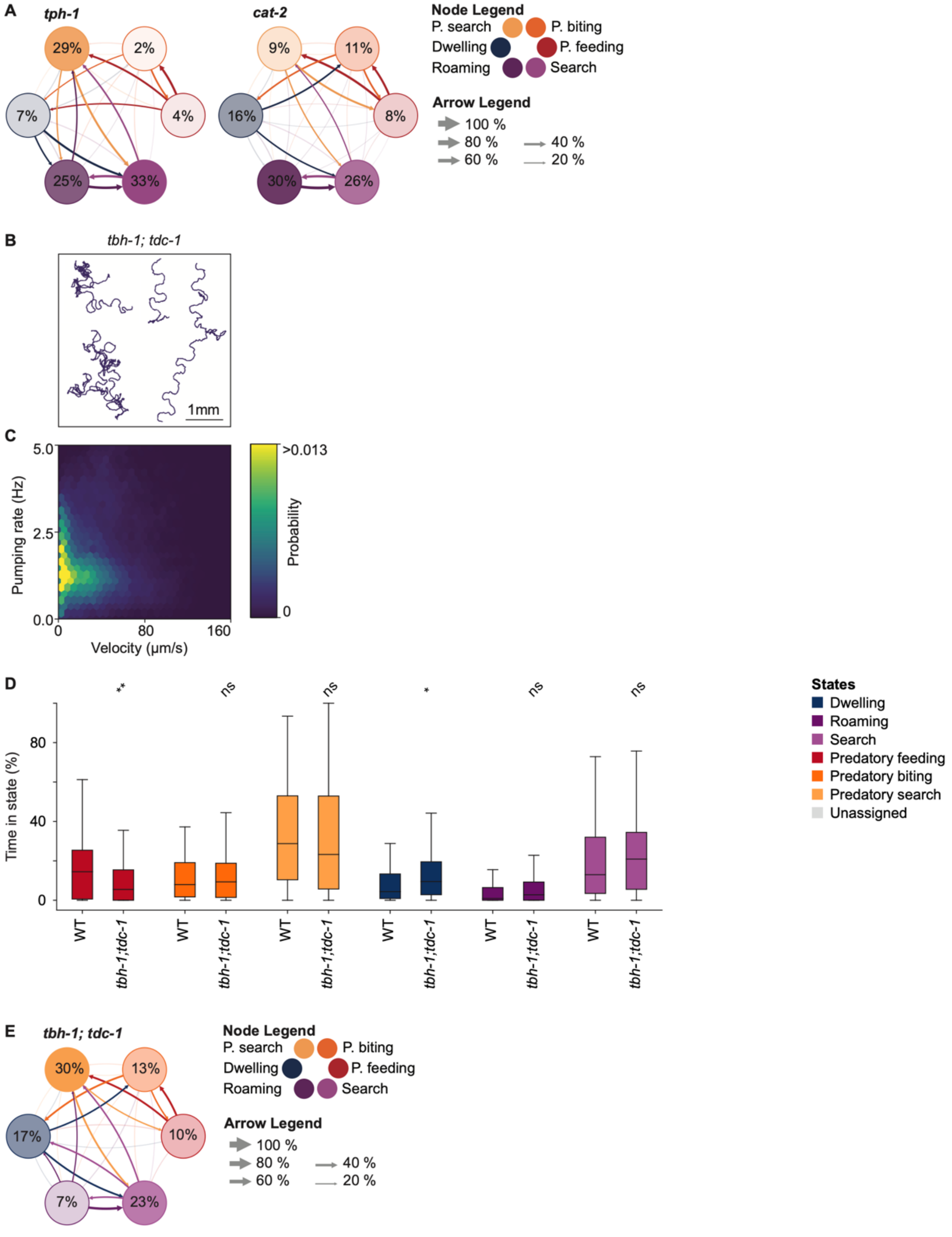

A
tph-1
cat-2
29%
2%
7%
4%
25%
33%
9%
11%
16%
8%
30%
26%
Node Legend
P. search
P. biting
Dwelling
P. feeding
Roaming
Search
Arrow Legend
100 %
80 %
60 %
40 %
20 %
B
tbh-1; tdc-1
1mm
C
5.0
2.5
0.0
Pumping rate (Hz)
0
80
160
Velocity (µm/s)
>0.013
Probability
0
D
**
ns
ns
*
ns
ns
Time in state (%)
80
40
0
WT
tbh-1;tdc-1
States
Dwelling
Roaming
Search
Predatory feeding
Predatory biting
Predatory search
Unassigned
E
tbh-1; tdc-1
30%
13%
17%
10%
7%
23%
Node Legend
P. search
P. biting
Dwelling
P. feeding
Roaming
Search
Arrow Legend
100 %
80 %
60 %
40 %
20 %

**Fig. S5. Behavioral state prediction for *tph-1, cat-2*, and *tbh-1; tdc-1***

(**A**) Behavioral state transitions for *tph-1* and *cat-2* mutants. The number in circles indicates the average state duration and the arrow size indicates the transition rate normalized to outgoing transitions. (**B**) Example tracks and (**C**) probability density map of velocity and pumping rate for the *tbh-1; tdc-1* double mutant. (**D**) Relative time in each behavioral state for WT versus the *tbh-1; tdc-1* double mutant. (**E**) Same as in (A) the average transition rates between behavioral states for the *tbh-1; tdc-1* double mutant.

Supp. 6

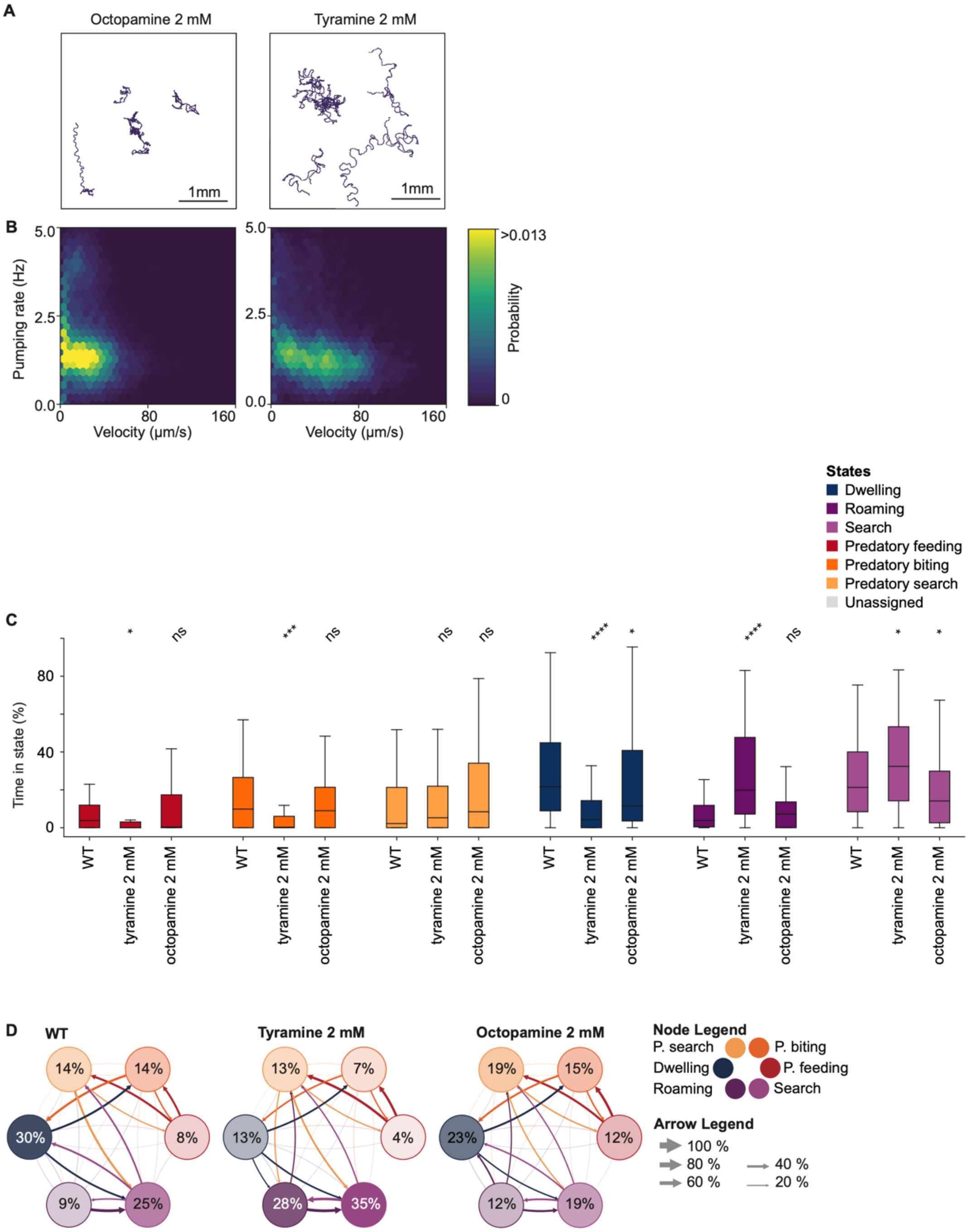
A
Octopamine 2 mM
Tyramine 2 mM
1mm
1mm
B
Pumping rate (Hz)
5.0
2.5
0.0
0
80
160
Velocity (µm/s)
>0.013
Probability
0
States
Dwelling
Roaming
Search
Predatory feeding
Predatory biting
Predatory search
Unassigned
C
Time in state (%)
0
40
80
* ns *** ns ns ns **** * **** ns * *
WT
tyramine 2 mM
octopamine 2 mM
D
WT
Tyramine 2 mM
Octopamine 2 mM
Node Legend
P. search
P. biting
Dwelling
P. feeding
Roaming
Search
Arrow Legend
100 %
80 %
60 %
40 %
20 %
14%
14%
30%
8%
9%
25%
13%
7%
13%
4%
28%
35%
19%
15%
23%
12%
12%
19%

**Fig. S6. Behavioral state prediction for exogenous application of octopamine and tyramine**
(**A**) Example tracks and (**B**) probability density map of velocity and pumping rate for animals exposed to 2 mM octopamine or tyramine, respectively. (**C**) Relative time in each behavioral state for WT compared to animals on 2 mM octopamine or tyramine. (**D**) Behavioral state transitions for WT, 2 mM tyramine, and 2 mM octopamine. The number in circles indicates the average state duration and the arrow size indicates the transition rate normalized to outgoing transitions.

Supp. 7

Ppa-ser-3

bnn115

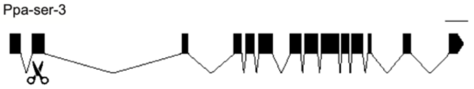

LVVIVVLGNSLVIAAVLLRRRLRSATGLLILSLGVADLLVGTVILPFSIANEVL
7bp del LVVIVVLGNSLVIASFAGGCDRPLACSFSPLEWPTY*

Ppa-ser-6

bnn116

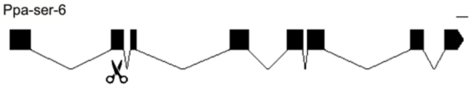

QEHHERWLYGEIFCRAWLSLDVFLSTASIYNLLAISVDRYMAVRQPIRYPIISSKRFVRQMIAG
16 bp del QEHHERYSVEHGYLLTYSFPLRVSIIYSRLVWIDTWQLDNQFDIPSFHLKDSSVR*

Ppa-octr-1

bnn119

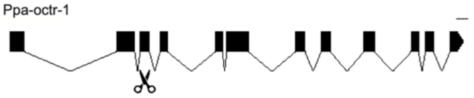

CEMWLALDVLFVTASILHICAISLDRYFCVTQPLSYPNKRTPLRMMLMIGCSWLVSLLISLPPIFGLRPPDRG
7bp ins CEMWLDGSTGCPLRDRVHPSHLCHFPGQVLLRNAAPFLSQQAHASQDDANDWLFLACISSDFATTNFRATTT*

Ppa-tyra-2

bnn123

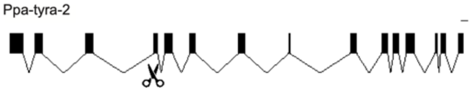

MAKRNKKTAGIMILSVWVSSALISLAPTLGWKQIADAPNMQMQNGTLQCTFLDLPSYTVYSATGSFFIPTLVMFFVYFKIYQAFAK
8bp del MAKRDGWHNDIIGVGVECTHLSGAHVGLEADRGCAQYADAERDITVYLPRSPFVHGVLCYGFLLHSHSRYVLRLLQNLPSVR*

Ppa-ser-2

bnn125

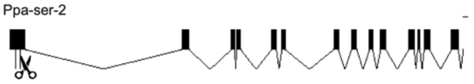

SALLTKGVEEEHTPKRESALLRPILPTHPTRPTPAAVMSAAPASSGAQPPTQDPMVLSQLFSITASIANSTTAIDVTPIPPPPSIIPIVELVLGTIT
7bp del SALLTKGESTHQKGNPPSSVRSFRPIRPVRHLQQSCLLHQHHPARSHQPKIRWCYRSSSPSQHRSPIRRRPSMSRQSRHRPPSFLSWNSCWALSHIW*

Ppa-lgc-55

bnn129

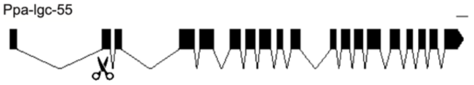

GIYCDLRSQYCKTQESGRQSCECRDGFVQADNGTCYMSRALANSMEAEPTEFCILGQAEKDATRILNDILKKYDKNLVP
4bp del GIYCDLRSQYCKTQVEDKVVNVEMDSYRQTMGHVTCLVHWRIRWKLNRRSSVFWDRLRKMQPEFSMIFSRNMIRISSQR*

Ppa-tyra-3

bnn132

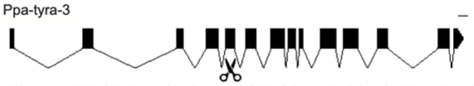

ITSPLKYPRTRSRKMAAGLLSAVWLLSFIVCSPPLFVQGWGHTPDDPPASQRESHITCSYSM
1bp del ITSPLKYPRTRSRKMAAGLSRPFGCFRSSFVLHRFSFKDGVIPPMIHQLHKENPTLLALIQ*

Ppa-tyra-2

bnn139

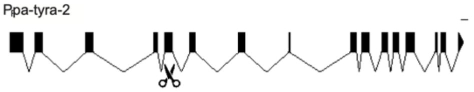

DYMAKRNKKTAGIMILSVWVSSALISLAPTLGWKQIADAPNMQMQNGTLQCTFLDLPSYTVYSATGSFFIPTLVMFFVYFKIYQA
10 bp sub DYMAKRNKKTAGIIGVGVECTHLSGAHVGLEADRGCAQYADAERDITVYLPRSPFVHGVLCYGFLLHSHSRYVLRLLQNLPSVR*

Ppa-ser-2

bnn138

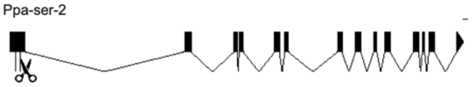

VVRAELSSLDPSALLTKGVEEEHTPKRESALLRPILPTHPTRPTPAAVMSAAPASSGAQPPTQDPMVLSQLFSITASIANSTTAIDVTPIPPPPSIIPIVELV
LGTITYLVIIAMTVVGNTLVVVAVFSYRPLKKVQNYLLVSLAASDLAVAIFVMPLHVVKFLADGKWLLGIWVCQFFTTSDILLCTSSILNLCAI

8 del VVRAELSSLDPSALLTKGGAHTKKGIRPPPSDPSDPSDPSDTCSSHVCCTSIIRRAATNPRSDGVIAALLHHSIDRQFDDGHRCHANPATALHHS
YRGTRVGHYHISGDHCDDRRGQYPRRCCRLLISTTQEGPKLSARLSGSIRSCRCHLCHAATRSQVSRRWQVAIGHMGLSVLHDFRYSSMHIFDPQSVCDCYR*

**Fig. S7. CRISPR/Cas9 mutants generated.**

Gene structures and protein sequence alignments of all mutants generated by CRIPSR/Cas9 in this study. Scale bar = 100 bp.

Supp. 8

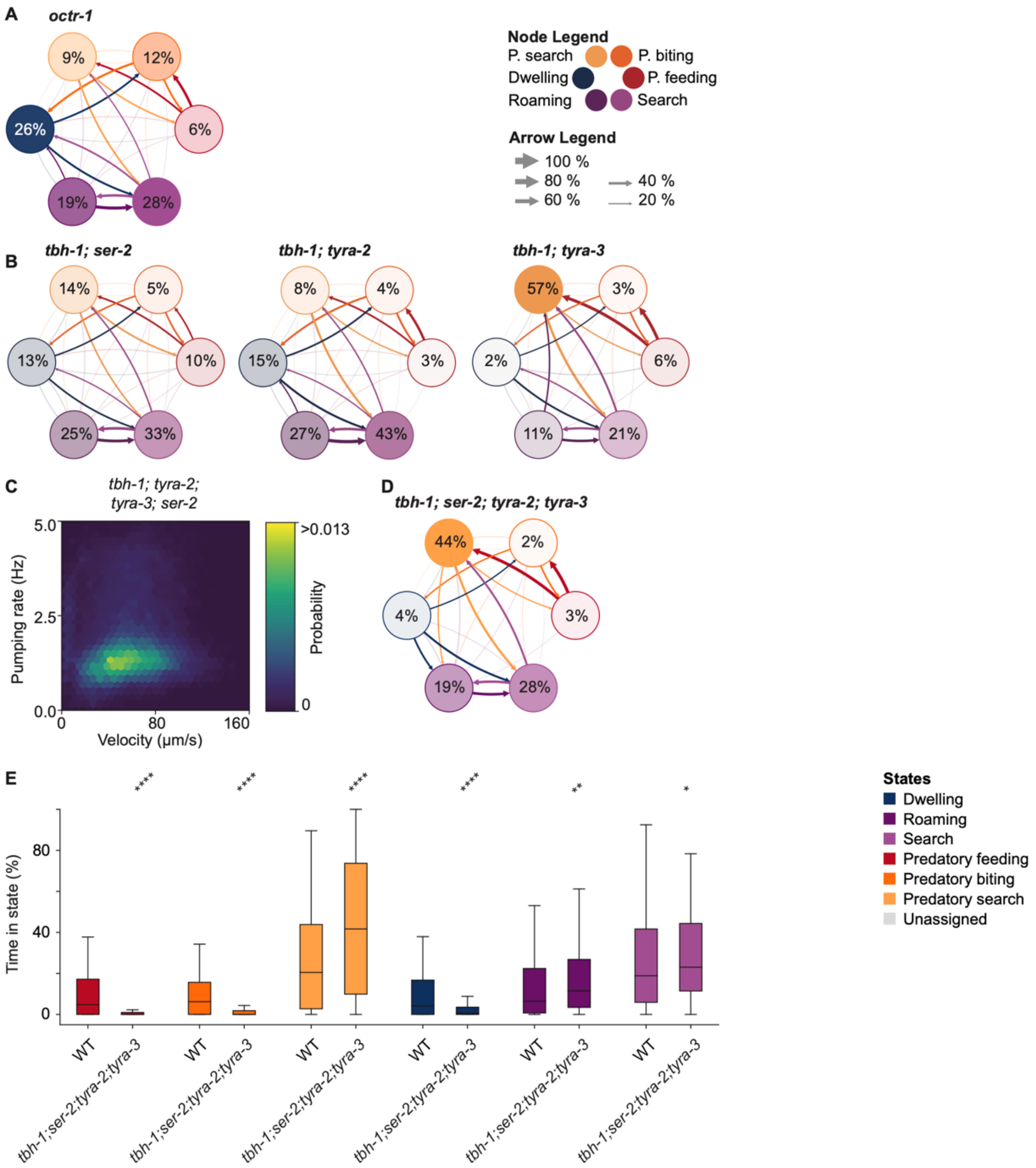
A
octr-1
9%
12%
26%
6%
19%
28%
Node Legend
P. search
P. biting
Dwelling
P. feeding
Roaming
Search
Arrow Legend
100 %
80 %
60 %
40 %
20 %
B
tbh-1; ser-2
14%
5%
13%
10%
25%
33%
tbh-1; tyra-2
8%
4%
15%
3%
27%
43%
tbh-1; tyra-3
57%
3%
2%
6%
11%
21%
C
tbh-1; tyra-2; tyra-3; ser-2
5.0
2.5
0.0
Pumping rate (Hz)
0
80
160
Velocity (μm/s)
>0.013
Probability
0
D
tbh-1; ser-2; tyra-2; tyra-3
44%
2%
4%
3%
19%
28%
E
****
****
****
****
**
*
80
40
0
Time in state (%)
WT
tbh-1;ser-2;tyra-2;tyra-3
States
Dwelling
Roaming
Search
Predatory feeding
Predatory biting
Predatory search
Unassigned

**Fig. S8. Behavioral state prediction for *tph-1, cat-2*, and *tbh-1; tdc-1***

(**A**) Average transition rates between behavioral states for *octr-1*. The number in circles indicates the average state duration and the arrow size indicates the transition rate normalized to outgoing transitions. (**B**) Same as (A) but for *tbh-1; ser-2, tbh-1; tyra-2,* and *tbh-1; tyra-3*. (**C**) Probability density map of velocity and pumping rate for *tbh-1;ser-2;tyra-2;tyra-3*. (**D**) Same as (A) for the quadruple mutant *tbh-1; ser-2; tyra-2; tyra-3*. (**E**) Relative time in each behavioral state for WT compared to *tbh-1; ser-2; tyra-2 ;tyra-3* quadruple mutant.

Supp. 9

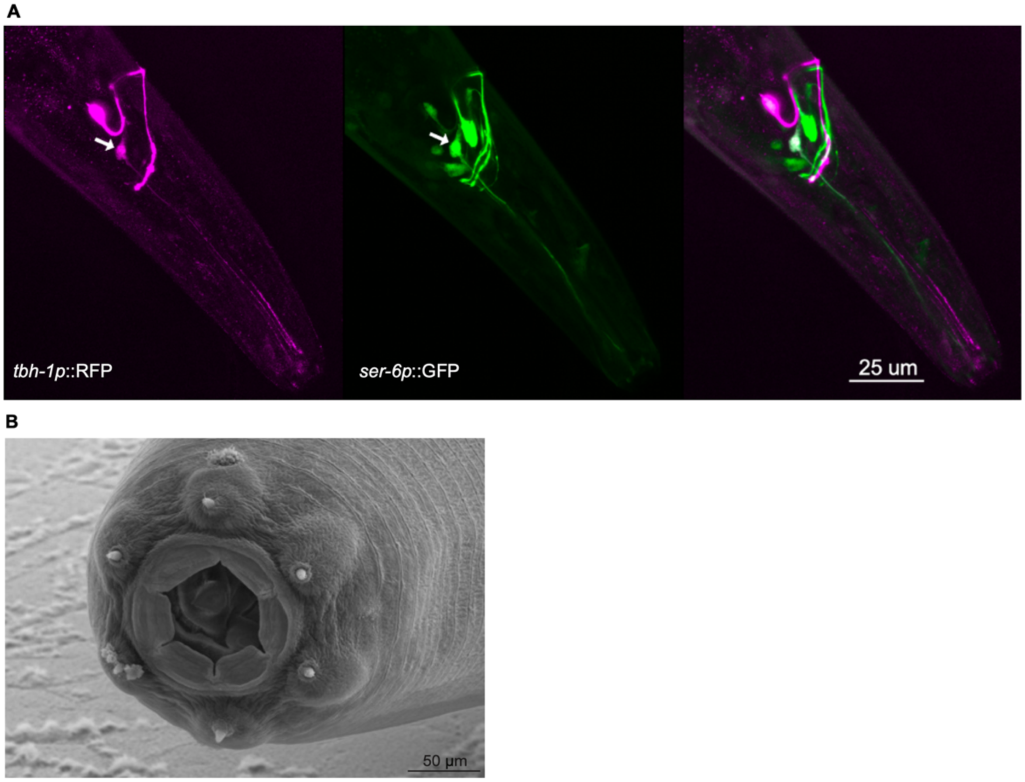

**Fig. S9. *P. pacificus* external sensory anatomy**

(**A**) Expression pattern of *tbh-1p*::RFP (magenta) and *ser-6p*::GFP (green) colocalization. Arrows indicate expression pattern overlap in a single neuron pair. Only left side of the animal shown for clarity. (**B**) SEM image of the *P. pacificus* face. Six sensory endings from the IL2 neurons circle the mouth opening and are candidates for prey detection.
